# Small-scale bioreactor cultivation of HEK293-based suspension cells increases extracellular vesicle yield

**DOI:** 10.64898/2026.07.14.738239

**Authors:** W. W. Woud, E. B. Dilla, N. Dits, T. Keijzer, C. Bernal, M. E. van Royen, E.S. Martens-Uzunova, J. de Vrij

## Abstract

**Purpose:** Extracellular vesicles (EVs) are increasingly explored as natural vehicles for drug delivery and gene therapy approaches. However, reproducible yield and scalability of EV production still pose major challenges in the clinical translation of EV-based therapies. In this study, we sought to quantify and characterize EVs released by suspension-cultured HEK293 cells (Expi293F cells) grown in shaker flasks or small-scale bioreactors, to investigate how the culturing environment affects EV production yield.

**Methods:** Expi293F cells were cultivated (N=3) in either shaker flasks or a bioreactor system, and total cell density, viability, and size were monitored. Supernatants were drawn daily post-cell seeding and were analyzed for EV quantity, size, morphology, and CD63 expression.

**Results:** No significant differences were observed in terms of total cell density, viability, and cell size between both cultivation settings. However, cultivation of Expi293F cells in the bioreactor environment significantly increased EV yield by 3-fold compared to shaker flask cultivation (p < 0.01). Other parameters such as average nanoparticle size, EV morphology, and CD63 expression remained comparable between both cultivation methods.

**Conclusion:** These results demonstrate that Expi293F-derived EV yield can be increased by culturing cells in a scalable bioreactor system. These findings pave the way towards the production of therapeutic-based EVs in a scalable and reproducible manner suitable for future (pre-)clinical applications.

## Introduction

Extracellular vesicles (EVs) are lipid bilayer membrane structures (30-8000 nm in diameter (Arraud et al. 2014)) released by cells. They are involved in cellular communication through transfer of surface receptors and/or a variety of macromolecules carried as cargo such as lipids, proteins, nucleic acids, protein-coding mRNAs and regulatory microRNAs (Pitt, Kroemer, and Zitvogel 2016; Simonsen 2017; Kumar et al. 2024). In nanomedicine, EVs are increasingly explored as natural vehicles for cell and gene therapies, and drug delivery applications. EVs have inherent targeting specificity, which allows them to be rapidly taken up by target cells (Kalra, Drummen, and Mathivanan 2016). Because of their endogenous nature EVs pose low cytotoxicity and immunogenicity in contrast to their synthetic and viral counterparts (Bader, Brigger, and Leroux 2024). Nevertheless, the yield and scalability of EV production still pose major challenges in the clinical translation of EV-based therapies.

Currently, therapeutic EVs are commonly obtained from human embryonic kidney (HEK293) cells. HEK293-based derivatives such as the suspension-grown Expi293FTM are currently investigated for their potential to produce (therapeutic) EVs in a controlled scalable manner (Sanz-Ros et al. 2023; Silva et al. 2021).

Large-scale cultivation of cell lines for various applications including the production of novel therapies requires the use of a bioreactor to provide optimal conditions to the cells (Stephenson and Grayson 2018). Whilst different types of bioreactors are employed in industrial manufacturing, scale down bioreactors are the preferred choice to perform small-scale screening studies in a cost and time efficient manner (Gallego-Murillo et al. 2022; Moreira et al. 2020). The possibility of running multiple experiments simultaneously allows for high throughput optimization and replication, which is essential for the statistical reproducibility of obtained results.

In the present study, a scaled down stirred-tank bioreactor system was assessed for its usability to reproducibly culture Expi293FTM cells and produce Expi293FTM-derived EVs. To this end, cells were cultivated in lab-scale shaker flasks or small-scale bioreactors, and growth parameters such as total cell density, viability, and cell size were analyzed. Supernatant samples from both cultivation settings were collected and analyzed for EV quantity, size, morphology, and expression of the tetraspanin CD63.

## Materials and Methods

### Bioreactor operation

Expi293™ cells (passage 13, 15, 17) (Gibco™) were cultivated in Expi293™ Expression Medium (Gibco™) in the 500 mL MiniBio reactor (Getinge) controlled by the my-Control system (Getinge), linked to Lucullus PIMS software (SecureCell) for data login and management. Per bioreactor run, the cells were seeded at a concentration of 0.5×106 cells/ml to a final volume of 300 ml. The dissolved oxygen (DO) and pH were measured by a LumiSens Optical DO probe (Getinge) and AppliSens pH sensor (Getinge), respectively. The temperature was kept at 37 °C using the pre-installed heating/cooling block and stirring speed was set at 300 rpm. The headspace was aerated with air (80% N2 and 20% O2). The setpoints and other settings used are summarized in Table 1.

**Table 1.** Bioreactor setpoints and other settings used during the cultivation of Expi293F cells.

|  |  |
| --- | --- |
| Bioreactor type | Applikon MiniBio 500 mL |
| Impeller type | Marine |
| Agitation speed | 300 rpm |
| pH setpoint | 7.10 (CO<sub>2</sub> sparged) |
| DO setpoint | 40% (Headspace aerated – 80% N<sub>2</sub> and 20% O<sub>2</sub>) |
| Sparging type | Open pipe |

### Shaker flask cultivation

To compare bioreactor EV production with EV production in a shaker flask environment, cells were also cultured in 125 ml Nalgene™ Single-Use PETG Erlenmeyer Flasks (Thermo Scientific™) at a working volume of 30 ml. Similarly, cells were seeded at a cell density of 0.5×106 cells/ml. The suspension cultures were maintained at 37 °C in the ISF1-X incubator (Thermo Scientific™) on a shaking platform (Thermo Scientific™ CO2 resistant shaker 8881102-19 mm orbit) at a frequency of 125 rpm.

### Cell culture sample drawing

A total culture volume of 5 mL was retrieved daily from the bioreactor through the sample port using a sterile HENKE-JECT® syringe with luer-lock tip (Henke Sass Wolf). Concurrently, 2 mL of crude culture was sampled daily from each shaker flask using a sterile serological pipette.

### Total cell concentration, viability, and size measurements

40 μL of cell culture sample was used to measure total cell density, viability, and cell size using the Countess 3 Automated Cell Counter (Thermo Fisher Scientific). For cell staining, the trypan blue exclusion test was performed in which raw cell culture, and 0.4% trypan blue were mixed in a 1:1 ratio. 10 µL of the resulting solution was pipetted into the chambers of the Countess™ slide (Invitrogen™).

### Clarified supernatants

Remaining cell culture samples were centrifuged for 5 mins at 300 x g to remove cell debris, and the resulting EV-containing clarified supernatants were collected and stored at -20 °C for EV quantification and characterization.

### Nanoparticle tracking analysis

Particle concentration in clarified supernatants was determined with nanoparticle tracking analysis (NTA) using the LM10 NanoSight (Nanosight Ltd, Salisbury, UK) equipped with a sCMOS camera and a blue 405-nm laser. Five individual videos of 60 seconds were recorded at camera level 12 and at a constant temperature of 20°C. Recordings were analyzed with a detection threshold set at 5, allowing discrimination of EVs from background particles. As a quality control, only measurements showing 40-90 particles per frame were considered.

### EVQuant

EVs in clarified supernatants were quantified with the EVQuant assay as described previously by Hartjes et al. (2020). Briefly, EVs in 40 µL clarified supernatant were fluorescently labeled with the generic Octadecyl Rhodamine B Chloride (Rhodamine-R18) membrane dye (Life Technologies). A Rhodamine-R18 dye-only control, a cell line (COLO205) derived EV isolate, and a 100-nm liposome sample were included to determine background levels and as internal standard controls. After background subtraction, the EV counts were converted to EV concentration.

### Transmission electron microscopy

EVs were visualized using transmission electron microscopy (TEM). For this, 10 μL droplets of clarified supernatant were deposited on formvar/carbon-coated 400 mesh Cu grids and incubated for 10 minutes. Afterwards, the remaining liquid was drained with a filter paper, and samples were stained with a drop of UranyLess stain for 1 minute. The remaining liquid was removed, and the grids were air dried. Grids were observed under the electron microscope Talos L120C TEM (Thermo Fisher Scientific) at 120 kV.

### Western Blot

Twenty-four µL of clarified supernatants were mixed with Laemmli sample buffer (non-reducing), and the resulting mixtures were heated at 95 °C for approximately 10 minutes. Afterwards, samples were loaded onto 10% SDS-PAGE gels (prepared in-house) and electrophoresed at 100 V for approximately 60 minutes. Subsequently, proteins were transferred onto Protran nitrocellulose membranes (Cytiva) and blocked (1h) at room temperature with 5% non-fat dry milk in Tris-Buffered Saline with 0.1% Tween-20. Blots were incubated overnight at 4°C with antibodies directed against CD63 (Clone: MX-49.129.5, Santa Cruz, cat#: sc-5275, 200 µg/ml, 1:200 dilution). Secondary antibodies (HRP-conjugated Goat anti-Mouse, 1:1000 dilutions, DakoCytomation) were incubated for 1h. BM Chemiluminescence Blotting Substrate (POD, Roche Life Science Products) was used to initiate the oxidation by HRP, and blots were imaged with the Amersham Imager 600 with a manual exposure of 12 minutes.

### Statistics

Statistical analysis was performed using R version 4.0.2 and Rstudio (Rstudio Team (2016). Rstudio: Integrated Development for R. Rstudio, Inc., Boston, MA; URL: http://www.rstudio.com/, version1.1.463). Statistical significance between EV concentrations and groups was determined using the standard student’s t-test, 95% CI with unpaired data. Coefficient of Variance (CV) scores were calculated for total cell density, viability, and cell size in samples drawn at the end of the cultivation process to analyze reproducibility between runs.

## Results

### Growth profiles of Expi293FTM cells

Analysis of total cell density, viability, and cell size revealed near-similar growth characteristics of Expi293™ cells cultivated in either bioreactors or shaker flasks across three independent runs (Figure 1. a-c). Doubling times of 20 to 24 hours were observed. High viability (>95%) and constant cell size (16.95 ± 0.19 μm, mean ± sd) were achieved throughout the cultivation period for both cultivation settings. Cell viability of cells cultivated in shaker flasks remained stable whereas bioreactor cultivated cells showed a marginal but statistically significant drop during the cultivation period (from 96.8% at day 0 down to 95.5% viability at day 4, p = 6.1E-4). Additionally, high reproducibility was achieved across the three independent process runs (CV scores <6%), indicating that cell growth in the shaker flask and the bioreactor is highly comparable. In summary, the growth characteristics of Expi293TM cells are maintained when cultured in either shaker flask or bioreactor environment, with the added value of larger working volumes and, in turn, increased cellular mass for bioreactor condition.

**Figure 1:**
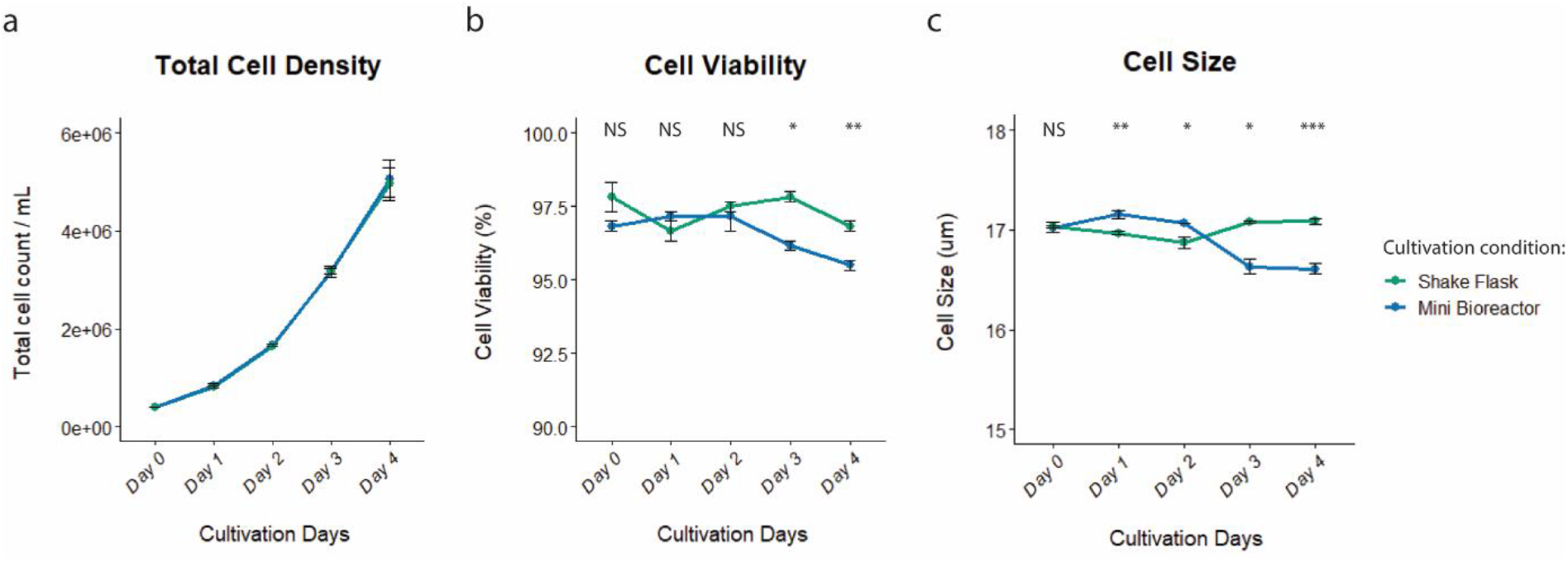
Comparison of Expi293F™ cell growth in shaker flasks and bioreactors. Three independent runs (n=3) were performed for each cultivation method and cell growth parameters were analyzed on clarified supernatants collected daily during the cultivation period. **a)** Analysis of total cell density indicated no differences between the cultivation settings. **b)** Analysis of cell viability and **c)** cell size indicated modest but statistically significant decreases for Expi293F cells cultivated in the bioreactors at the end of the cultivation period compared to cells cultivated in shaker flask conditions. Data shown as Mean ± SD for the three independent runs.

### EV analyses by NTA, EVQuant, TEM and Western Blot

Next, the EV production yield was compared between both cultivation settings. Clarified supernatants (collected during the cultivation period) were analyzed for their EV concentrations using NTA and EVQuant (Figure 2. a,b). Both techniques indicated a statistically significant (p < 0.05) increased EV yield after cultivation in the bioreactor compared to shaker flask cultivation, as measured at the end of the cultivation period (~3-fold and ~2-fold difference as measured by NTA and EVQuant, respectively). It should be noted that NTA detects all nanoparticles in suspension whereas EVQuant relies on the detection of fluorescent signals associated with lipid-based particles and hence is more specific for membrane particles. Using the EV concentrations as determined by EVQuant for samples obtained at the end of the cultivation period (Day 4), we calculated average EV-per-cell ratios of 8.14E^3^ and 1.73E^4^ for shaker flask and bioreactor culturing conditions, respectively. In other words, a 2-fold difference in EV production efficiency was demonstrated for Expi293F cells cultivated under bioreactor conditions compared to shaker flask conditions. Nanoparticle size analysis by NTA showed slight differences (non-significant) in terms of particle diameter between both cultivation settings (mean: 159.6 ± 2.8 nm vs 149.6 ± 1.1 nm for shaker flask and bioreactor, respectively - Figure 2. c). Visual examination of the clarified supernatant samples by TEM demonstrated the classical cup-shaped morphology of EVs (Figure 2. d), and WB directed against the putative exosome marker CD63 demonstrated increasing signal over the course of the cultivation period for either cultivation setting, with the signal being most pronounced for the bioreactor samples (Figure 2. e).

**Figure 2:**
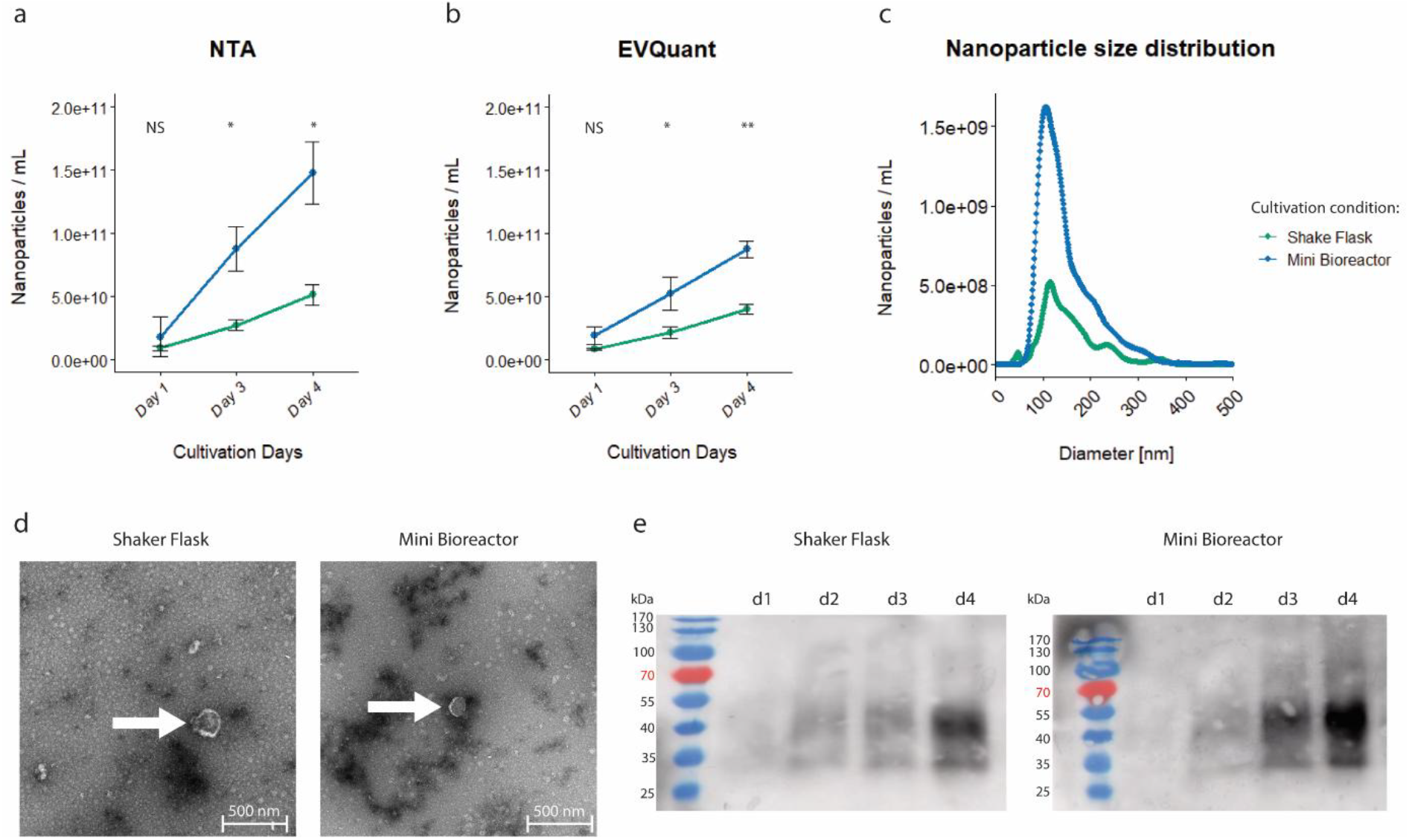
Comparison of Expi293F™ EV production in shaker flasks and bioreactors. Nanoparticle quantification by **a)** nanoparticle tracking analysis (NTA) and **b)** EVQuant demonstrated significantly increased nanoparticle yields by Expi293F™ cells cultivated in bioreactors compared to the shaker flask environment at the end of the cultivation period (NS: non-significant, *: p < 0.05, **: p < 0.01). **c)** Size analysis by NTA demonstrated similar profiles in samples drawn from either cultivation condition. Data shown represents measurements from one shaker flask (green) and one bioreactor (blue) sample. **d)** Morphological assessment of the nanoparticles revealed cup-shaped structures, indicative of EVs. **e)** Western blot directed against the putative exosome marker CD63 demonstrated increasing signal over the course of the cultivation period, with the strongest signals observed in the clarified supernatant samples drawn from the bioreactors. Data shown as Mean ± SD for the three independent runs.

## Discussion

Achieving the optimal balance between proper cell growth and high EV production yield is essential for the establishment of scalable cultivation methods suitable for large-scale therapeutic EV production. The current work shows the utility of bioreactors for the cultivation of Expi293FTM cells and demonstrates the successful production of EVs. Whilst total production cell concentrations were highly comparable between both cultivation settings, cell viability and cell size showed a modest but statistically significant difference for cells cultivated in bioreactors compared to shaker flasks (~95.5% vs 96.8%, and 16.6 vs 17.08 µm, respectively). However, we observed significantly increased EV production yield in the bioreactor set-up compared to the EV yield obtained in the shaker flask setting as demonstrated with both NTA and EVQuant. Cell viability in the bioreactor conditions was only marginally decreased at the end of the cultivation period, suggesting that the observed EV yield increase is not solely due to cell viability-related secretion of EVs (e.g. apoptotic bodies). Furthermore, no differences in terms of EV characteristics such as their mean size, morphology, and CD63 expression were observed between the cultivation conditions. These findings demonstrate the usability of scalable bioreactor systems for the production of EVs and pave the way to their utilization as therapeutic modalities.

## Statements and Declarations

### Author Contributions

All authors contributed to the study conception and design. The first draft of the manuscript was written by W.W. Woud and all authors commented on previous versions of the manuscript. All authors read and approved the final manuscript.

### Data Availability

The datasets generated during the study are available from the corresponding author upon request.

